# TSC22D4 interacts with Akt1 in response to metabolic and stress signals

**DOI:** 10.1101/2021.12.20.473554

**Authors:** Sevgican Demir, Gretchen Wolff, Annika Wieder, Adriano Maida, Marco Rahm, Martina Schnölzer, Stefanie Hauck, Julia Szendrödi, Stephan Herzig, Bilgen Ekim Üstünel

## Abstract

Transforming Growth Factor β 1 Stimulated Clone 22 D4 (TSC22D4) is an intrinsically disordered protein that regulates cellular and physiological processes such as cell proliferation, cellular senescence as well as hepatic glucose and lipid metabolism. The molecular mechanism of TSC22D4 action in these cellular and metabolic functions, however, remains largely elusive. Here, we identified TSC22D4 as a novel protein kinase B/Akt1 interacting protein, a critical mediator of insulin/PI3K signaling pathway implicated in diverse set of diseases including type 2 diabetes, obesity and cancer. TSC22D4 interacts with Akt1 not constitutively but rather in a regulatory manner. While glucose and insulin stimulation of cells or refeeding of mice impair the hepatic TSC22D4-Akt1 interaction, inhibition of mitochondria and oxidative stress, promote it; indicating that extra- and intra-cellular cues play a key role in controlling TSC22D4-Akt1 interaction. Our results also demonstrate that together with its dimerization domain, i.e. the TSC box, TSC22D4 requires its intrinsically disordered region (D2 domain) to interact with Akt1. To understand regulation of TSC22D4 function further, we employed tandem mass spectrometry and identified 15 novel phosphorylation sites on TSC22D4. Similar to TSC22D4-Akt1 interaction, TSC22D4 phosphorylation also responds to environmental signals such as starvation, mitochondrial inhibition and oxidative stress. Interestingly, 6 out of the 15 novel phosphorylation sites lie within the TSC22D4 D2 domain, which is required for TSC22D4-Akt1 interaction. Characterization of the regulation and function of these novel phosphorylation sites, in the future, will shed light on our understanding of the role of TSC22D4-Akt1 interaction in both cell biological and physiological functions. Overall, our findings postulate a model whereby TSC22D4 acts as an environmental sensor and interacts with Akt1 to regulate cell proliferation, cellular senescence as well as maintain metabolic homeostasis.

## Introduction

Insulin signaling pathway is an evolutionarily conserved mechanism that plays a key function in regulation of cell growth, cell proliferation, development, longevity, as well as metabolic homeostasis. When insulin binds to the insulin receptor (IR), IR gets activated and phosphorylates insulin receptor substrate (IRS) to initiate a cascade of phosphorylation events leading to activation of the insulin/phosphatidyl inositol 3-kinase (PI3K) signaling mediated by Protein Kinase B (PKB)/Akt [1]. Once phosphorylated by PI3K-dependent kinase 1 (PDK1) and mechanistic target of rapamycin complex 2 (mTORC2) on T308 and S473 respectively, Akt kinase becomes catalytically active and phosphorylates a wide range of targets including glycogen synthase kinase 3β (GSK3β), forkhead box O1 (FoxO1), tuberous sclerosis complex 2 (TSC2) to promote cell survival, cell growth and cell proliferation, as well as to regulate glucose and lipid metabolism [2]. In addition to the phosphorylation events, protein-protein interactions play a key role in regulating Akt function [3]. PtdIns-3,4,5-P3 (PIP3), c-jun N terminal kinase interacting protein 1 (JIP1), growth factor receptor-binding protein 10 (Grb10), Tribbles homolog 3 (Trb3) represent only a few of the proteins that interact with Akt to regulate its distinct functions [3–5].

Elevated insulin levels during overnutrition initiates negative feedback loops in insulin signaling pathway, leading to the pathological condition known as insulin resistance, in which metabolic organs fail to respond to circulating insulin levels. Overnutrition also induces a chronic low-grade inflammation which activates c-jun N terminal kinase (JNK) and inhibits AMP-activated protein kinase (AMPK), contributing to pathogenesis of insulin resistance. When necessary measures such as promoting weight loss via healthy eating and physical exercise are not taken, insulin resistance may progress further into type 2 diabetes, which is hardly reversible and leads to several complications in peripheral organs in the long run [6].

Recently, we identified Transforming Growth Factor β1 (TGFβ1) Stimulated Clone 22 D4 (TSC22D4) as a regulator of Akt signaling pathway in mouse models of type 2 diabetes [7]. TSC22D4 belongs to TSC22 protein family, which contains the evolutionarily conserved TSC box with a leucine zipper motif. TSC22 family consists of 4 proteins: TSC22D1, TSC22D2, TSC22D3 and TSC22D4 with alternatively spliced isoforms and they can homo- and heterodimerize with each other via leucine zipper motif in TSC Box to regulate cell biological functions such as cellular senescence, cell proliferation and apoptosis [8–13].

Despite discovered two decades ago, our current knowledge on the regulation and function of TSC22D4 remains very limited. TSC22D4 expression increases in cultured kidney cells upon osmotic stress and TSC22D4 subcellular localization in neurons alters during stages of the embryonic development and differentiation [10, 13]. TSC22D4 also suppresses cellular senescence by impairing junB function [14]. Previously, we have shown that hepatic TSC22D4 expression increases upon liver damage and cancer cachexia, and reprograms liver lipid metabolism [15]. Targeting TSC22D4 expression in the livers of mice with type 2 diabetes improves insulin-induced Akt phosphorylation, alleviates insulin resistance and hyperglycemia [7]. Strikingly, in obese human patients, elevated TSC22D4 expression in the liver positively correlates with insulin resistance [7]. These studies clearly establish a novel and reciprocal connection between TSC22D4 and metabolic regulation, yet the molecular mechanisms of this connection remains largely elusive.

Here, we identify TSC22D4 as a novel Akt1 interacting protein. Glucose stimulation of the starved cells destabilizes the TSC22D4-Akt1 interaction, which is exacerbated further upon insulin stimulation. Mitochondrial inhibition and oxidative stress, on the other hand, promote the TSC22D4-Akt1 interaction. To understand TSC22D4 regulation further, we employed liquid chromatography tandem mass spectrometry (LC-MS/MS) and identified 15 novel phosphorylation sites on TSC22D4. Similar to TSC22D4-Akt1 interaction, TSC22D4 phosphorylation also responds to both metabolic and stress signals suggesting that TSC22D4 may act as a hub where it receives signals from different upstream kinases to control intracellular crosstalk events. Overall, based on our findings, we propose a model in which TSC22D4 acts as a signaling molecule that senses the environmental cues to interact with and regulate Akt1 function.

## Results

### TSC22D4 interacts with Akt1

In our previous studies, we showed that acute TSC22D4 knockdown promotes insulin induced Akt phosphorylation both in cultured primary hepatocytes and in mouse livers. Interestingly, TSC22D4 knockdown also improves glucose handling and insulin sensitivity in diabetic mouse models [7]. To study the function of TSC22D4 further, we generated hepatocyte specific TSC22D4 knockout mice (TSC22D4^hep-/-^) (Supp Fig. 1A). Similar to our previous findings, TSC22D4^hep-/-^ mice had improved glucose handling in glucose tolerance test and lower fasting blood glucose levels compared to TSC22D4^flox/flox^ control mice when challenged with high fat diet without any changes in body weight (Supp Fig. 1B-D). In primary hepatocytes isolated from TSC22D4^hep-/-^ and TSC22D4^flox/flox^ control mice, we observed that TSC22D4 knockout cells had a much higher insulin induced phosphorylation of Akt and its direct downstream targets GSK3β and FoxO1 as well phosphorylation of its indirect target S6K1, which agree with our previous studies performed with shRNA-mediated TSC22D4 knockdown in primary hepatocytes and livers (Supp Fig. 1E) [7].

To understand the molecular mechanisms of TSC22D4 action on insulin/Akt signaling pathway, we examined the functional domains on TSC22D4 protein in detail. Interestingly, TSC22D4 does not have any domains with known function other than its TSC box that contains a leucine zipper motif (Fig. 1A). TSC box is common to all TSC22 family members and plays role in homodimerization or hetero-dimerization of TSC22 family members with each other [8, 16]. By performing Clustal-omega analysis, we identified two other regions (Region 1 and Region 2: R1 and R2) at TSC22D4 N-terminus which are highly conserved from fish to mammals (Fig. 1A and Supp. Fig. 2) [17]. The primary sequence flanked by R2 and TSC Box (105-308 aa) is also highly conserved between humans and mice, yet does not contain any particular three-dimensional structure such as alpha-helices or beta sheets, hence fall into the category of instrinsically disordered region (Fig. 1A and Supp. Fig. 2). Until recently, intrinsically disordered regions were mainly ignored due to lack of a defined three-dimensional structure, yet these regions contain many different post-translational modification (PTM) sites [18]. This diversity and multiplicity of PTMs within intrinsically disordered regions allow proteins to acts as critical sensors of the environmental cues as well as enable them to act as hubs for crosstlaking between signaling pathways.

**Figure 1.**
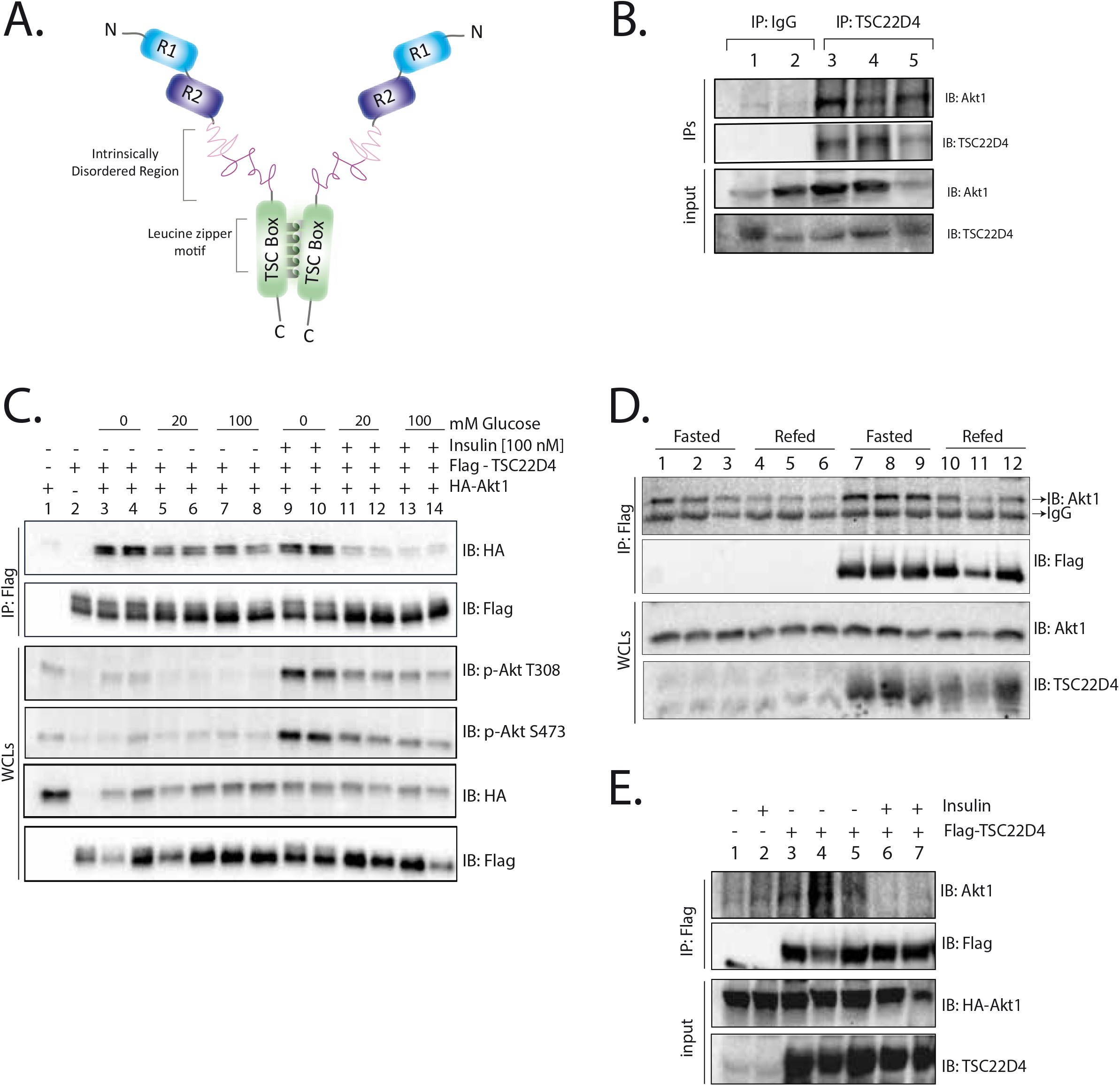
TSC22D4 interacts with Akt1. **A**. Illustration of TSC22D4 protein with evolutionarily conserved domains. **B.** Endogenous TSC22D4 from the wild type mouse liver lysates was immunoprecipitated (IP) with TSC22D4 antibody and the IPs and liver lysates (input) were immunoblotted (IB) with indicated antibodies. Normal rabbit IgG antibody was used as negative control to TSC22D4 antibody. **C.** Hepa 1-6 cells were transiently transfected with vector control or co-transfected with Flag-TSC22D4 (2,5 μg) and HA-Akt1 (2,5 μg) plasmids. 30 h posttransfection cells were serum and glucose starved overnight. Cells were pretreated without or with glucose [20 mM] for 30 min and incubated in the absence or presence of insulin [100 nM] for an additional 30 min and then lysed. Flag-TSC22D4 was immunopecipitated (IP) with anti-Flag affinity gel and the IPs and whole cell lysates (WCL) were immunblotted (IB) with indicated antibodies. **D.** Flag-TSC22D4 was IPed from the liver lysates of mice that were injected with adenoviruses containing empty vector control or Flag-TSC22D4 cDNA. The mice were starved for 16 h and refed the next day prior to sacrifice. The IPs and WCLs were IBed with indicated antibodies. **E.** Flag-TSC22D4 was immunoprecipitated (IP) with Flag affinity gel from liver lysates of 6 h starved and insulin injected mice (1.5U/kg for 10 min). IPs and whole liver lysates (input) were immunoblotted with indicated antibodies.

Since TSC22D4 contains a relatively long stretch of an intrinsically disordered region, we asked whether TSC22D4 interacts with Akt to regulate its function. We prepared whole liver lysates from wild type mice and immunoprecipitated endogenous TSC22D4. In western blots, we showed that endogenous Akt1 was indeed enriched in endogenous TSC22D4 co-immunoprecipitates compared to IgG controls, demonstrating that hepatic TSC22D4 and Akt1 interact *in vivo* in mouse livers (Fig. 1B). Unlike Akt1, Akt2 levels in the TSC22D4 co-immunoprecipitates failed to enrich over IgG controls, suggesting that TSC22D4 specifically interacts with Akt1 rather than Akt2 (data not shown).

In order to study the regulation of TSC22D4-Akt1 interaction more in detail, we employed Hepa 1-6 cells. We transiently transfected Hepa 1-6 cells with Flag-TSC22D4 and HA-Akt1 plasmids and repeated the co-IP experiments. As shown in Fig. 1C, HA-Akt1 co-IPed with Flag-TSC22D4 and interestingly glucose stimulation impaired the TSC22D4-Akt1 interaction. Increasing glucose concentration from 20 mM to 100 mM did not impair the TSC22D4-Akt1 interaction any further, but additional stimulation of cells with 100 nM of insulin did (Fig. 1C, compare lanes 5-8 to 11-14). Very interestingly, in the absence of glucose, insulin alone failed to disrupt the TSC22D4-Akt1 interaction (Fig. 1C, compare lanes 9-10 to 11-14). Similar to glucose and insulin stimulation in the cells, refeeding of wild type mice after an overnight starvation or i.p. insulin injection impaired the hepatic TSC22D4-Akt1 interaction (Fig. 1D, 1E). Overall these data indicate that TSC22D4 is a novel Akt1 interacting protein and metabolic signals such as glucose and insulin stimulation impair the TSC22D4-Akt1 interaction.

### Stress signals promote TSC22D4-Akt1 interaction

Both in mice and in the cells, the TSC22D4-Akt1 interaction was strongest during starvation and was impaired upon glucose and insulin stimulation. Since glucose availability promotes ATP generation, we tested whether mitochondrial function regulates the TSC22D4-Akt1 interaction as well. We performed Flag-TSC22D4 and HA-Akt1 co-IPs upon pharmacological inhibition of mitochondria with complex I and III inhibitors, rotenone and antimycin, respectively. As shown in Fig. 2A, rotenone and antimycin co-treatment (lanes 6-7) promoted the TSC22D4-Akt1 interaction and reduced Akt-S473 phosphorylation while increasing AMPK-T172 phosphorylation confirming the increase in AMP/ATP ratio upon drug treatments. Next, we asked whether inhibition of mitochondria promotes TSC22D4-Akt1 interaction via AMPK activity. To this end, we employed U2OS-WT and U2OS-AMPKα1/α2 double knockout (DKO) cells to transiently transfect Flag-TSC22D4 and HA-Akt1 plasmids and repeated co-IP experiments in the presence or absence of rotenone/antimycin. As shown in Fig. 2B, lack of AMPK did not affect rotenone/antimycin induced TSC22D4-Akt interaction. Additionally, glucose and insulin stimulation impaired TSC22D4-Akt interaction to similar degree in the presence or absence of AMPK (Fig. 2C). Overall these data suggest that glucose stimulation or inhibition of mitochondria regulate TSC22D4-Akt1 interaction independent of AMPK signaling.

**Figure 2.**
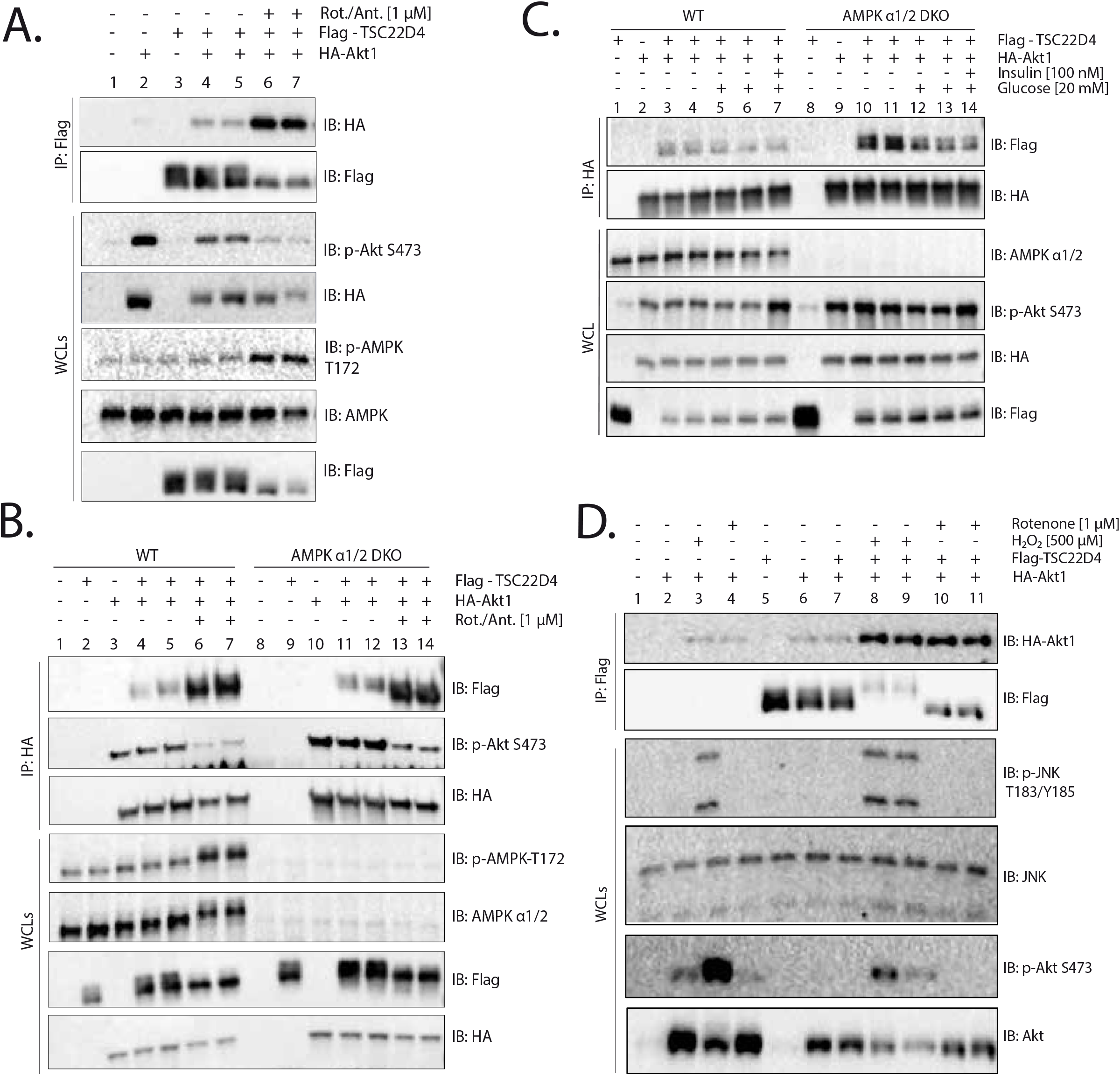
Stress signals promote TSC22D4-Akt1 interaction. **A.** Hepa 1-6 cells were transiently transfected with vector control or co-transfected with Flag-TSC22D4 (2,5 μg) and HA-Akt1 (2,5 μg) plasmids. 30 h posttransfection cells were serum and glucose starved overnight. Cells were pretreated without or with rotenone [1μM] and antimycin [1μM] mix 1 h prior to lysis. Flag-TSC22D4 was immunopecipitated (IP) with anti-Flag affinity gel and the IPs and whole cell lysates (WCL) were immunblotted (IB) with indicated antibodies. **B.** Similar to A except that WT or AMPKα1/2 Double Knockout U2OS cells were used and cells were starved for 4-6 h. IPs were performed with anti-HA antibody. **C.** WT and AMPKα1/2 Double Knockout U2OS cells were transiently transfected with vector control or co-transfected with Flag-TSC22D4 (2,5 μg) and HA-Akt1 (2,5 μg) plasmids. 48 h posttransfection cells were serum and glucose starved for 4-6 h. Cells were pretreated without or with glucose [10 mM] for 30 minutes and incubated in the absence or presence of insulin [100 nM] for an additional 30 min and then lysed. HA-Akt1 was immunopecipitated (IP) with anti-HA antibody and the IPs and whole cell lysates (WCL) were immunblotted (IB) with indicated antibodies. **D.** Hepa 1-6 cells were transiently transfected with vector control or Flag-TSC22D4 (2,5 μg) and HA-Akt1 (2,5 μg) plasmids. 30 h posttransfection cells were serum and glucose starved overnight. Cells were pretreated without or with H_2_O_2_ [500 μM] or Rotenone [1 μM] for an additional 1 h and then lysed. Flag-TSC22D4 was immunopecipitated (IP) with anti-Flag affinity gel and the IPs and WCLs were immunblotted (IB) with indicated antibodies.

Mitochondrial dysfunction contributes to generation of reactive oxygen species and induces oxidative stress. Hence, we asked whether treatment of cells with ROS would also have an effect on TSC22D4-Akt1 interaction and indeed found out that H_2_O_2_ treatment of Hepa 1-6 cells robustly promoted TSC22D4-Akt1 interaction (Fig. 2D). H_2_O_2_ treatment also induced JNK phosphorylation indicating the activation of stress signals. Therefore, we also tested whether JNK directly regulates TSC22D4-Akt1 interaction. Interestingly, TSC22D4-Akt1 interaction remained completely intact in JNK1/2 DKO mouse embryonic fibroblasts (MEF) demonstrating that TSC22D4-Akt1 interaction takes place independent of JNK1/2 (data not shown). Overall, these data show that while anabolic signals such as glucose and insulin stimulation impair the TSC22D4-Akt1 interaction, catabolic and stress signals promote it, independent of AMPK and JNK signaling pathways, respectively.

### TSC22D4-Akt interaction requires TSC22D4 intrinsically disordered region

To map the TSC22D4 domain(s) required for TSC22D4-Akt interaction, we created TSC22D4 deletion mutants as depicted in Fig. 3A. Since the intrinsically disordered region region contains a relatively long stretch of aminoacids, we arbitrarily splitted it into two as Domain 1 (D1) and Domain 2 (D2) (Fig. 3A). We transiently co-transfected Hepa 1-6 cells with different Flag-TSC22D4 deletion mutants and HA-Akt1. Deletion of R1 (ΔR1) and R2 (ΔR2) individually or together (ΔN) as well as deletion of the first half of intrinsically disordered region i.e. ΔD1, did not impair TSC22D4’s ability to interact with HA-Akt1 although ΔR2- and ΔN-TSC22D4 also expressed at very low levels compared to other TSC22D4 alleles (Supp. Fig. 3A).

**Figure 3.**
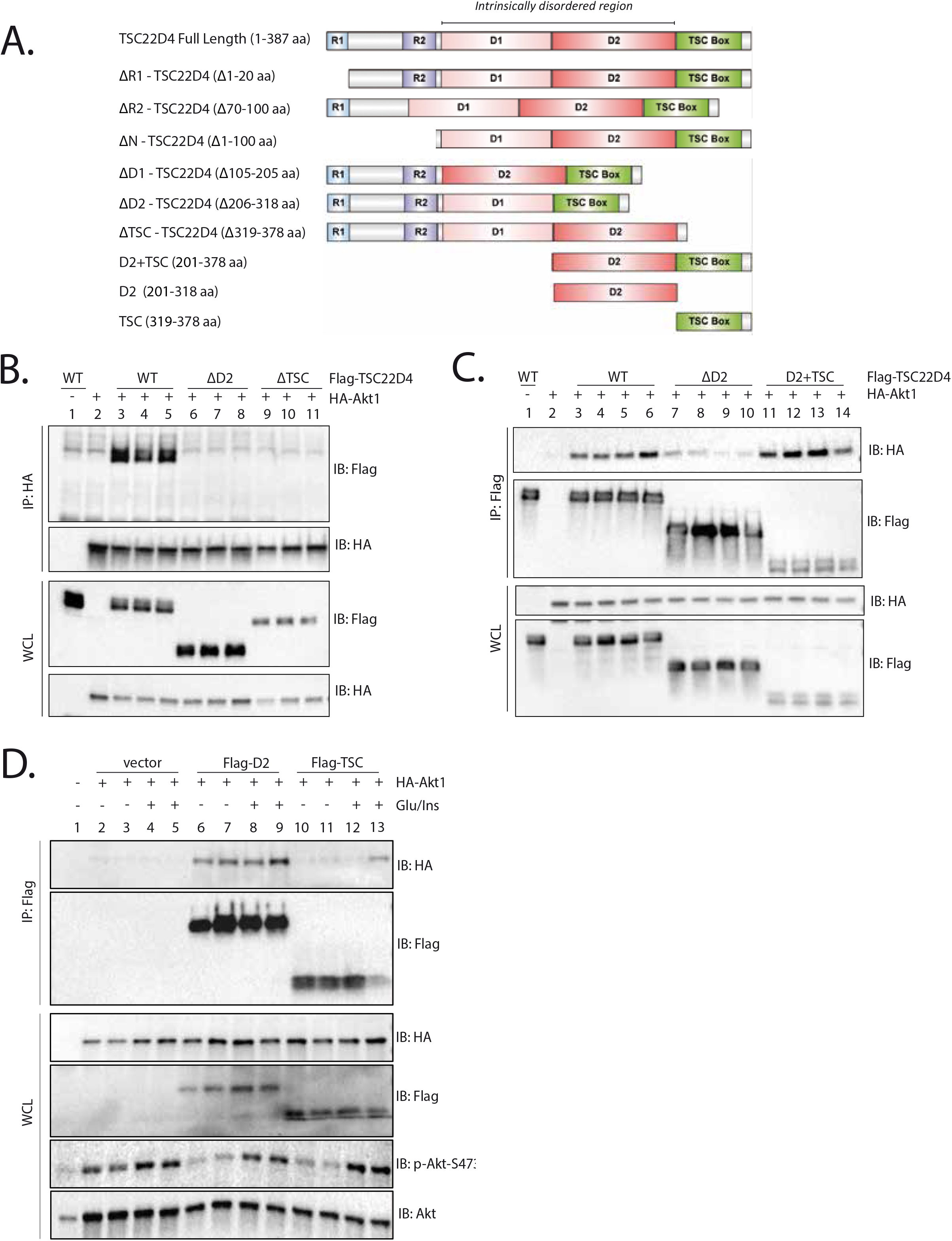
D2 Domain and TSC Box are required for TSC22D4-Akt interaction. **A**. Illustration of full length TSC22D4 and deletion mutants we generated. **B.** Hepa1-6 cells were transiently co-transfected with Flag-TSC22D4-WT (2,5 μg), -ΔD2 (2,5 μg) or -ΔTSC (2,5 μg) deletion mutants and HA-Akt1(2,5 μg). HA-Akt1 was immunopecipitated (IP) with anti-HA antibody and the IPs and whole cell lysates (WCL) were immunblotted (IB) with indicated antibodies. **C.** Hepa1-6 cells transiently co-transfected with Flag-TSC22D4-WT (2,5 μg), -ΔD2 (2,5 μg) or D2+TSC (2,5 μg) alleles and HA-Akt1 (2,5 μg). Flag-TSC22D4 was immunoprecipitated (IP) with anti-Flag affinity gel and the IPs and WCL were immunblotted (IB) with indicated antibodies. **D.** Hepa1-6 cells transiently co-transfected with Flag-tagged empty vector control (4 μg), D2 domain (4 μg) or TSC Box (4 μg) truncation mutants and HA-Akt1(1 μg). 30 h post-transfection, cells were serum and glucose starved overnight, and the next day they were stimulated without or with glucose [20 mM] for 30 min followed by insulin [100 nM] stimulation for another 30 min prior to lysis. Flag IPs and WCLs were immunoblotted (IB) with indicated antibodies.

Unlike R1, R2 and D1 regions, deletion of D2 (ΔD2) or TSC box (ΔTSC), however, completely attenuated the TSC22D4-Akt interaction (Fig. 3B). Even when the cells were treated with rotenone and antimycin that boost the TSC22D4-Akt1 interaction, the ΔD2-TSC22D4 mutant failed to interact with Akt completely (Supp. Fig. 3B). The D2 domain lies within the intrinsically disordered region and potentially carries the necessary features of undergoing versatile PTM events to interact with different partner proteins similar to other intrinsically disordered region proteins. The TSC box, however, plays a key role in homodimerization which might also be critical for the TSC22D4-Akt1 interaction. Indeed the Flag-TSC22D4-ΔTSC mutant failed to dimerize with full length myc-TSC22D4, confirming the requirement of TSC box for TSC22D4 homodimerization (Supp. Fig. 3C). The ΔD2-TSC22D4, on the other hand, still managed to homodimerize with myc-TSC22D4, albeit to a less extent compared to wild type Flag-TSC22D4 (Supp. Fig. 3D). Overall, these data suggest that TSC domain might have an indirect role in mediating TSC22D4-Akt1 interaction via homodimerization.

Next, we generated the D2+TSC mutant to address whether it will be sufficient to interact with Akt1. Indeed the D2+TSC allele interacted with Akt1 at stronger levels compared to WT-TSC22D4 (Fig. 3A, Fig. 3C, Supp Fig. 3E).Next, we created the D2 domain and TSC box alone truncated mutants and repeated the co-IP experiments. Interestingly, while D2 domain alone succesfully interacted with Akt1, the TSC box alone failed to do so (Fig. 3D). Overall, these data show that although both D2 domain and TSC box are required to interact with Akt, only D2 domain is sufficient to maintain the interaction.

### TSC22D4 undergoes phosphorylation

During experiments in which we inhibitied mitochondria with rotenone and antimycin, we observed that the Flag-TSC22D4 signal collapsed to a one single band at a lower molecular weight in the western blots, indicating the loss of certain PTMs (Fig. 2A, 2B). Similar to the overexpressed Flag-TSC22D4, endogenous TSC22D4 also behaved in a similar manner, and lost PTMs upon overnight starvation both in Hepa 1-6 and AML12 cells (Fig. 4A and B). As opposed to rotenone and antimycin, H_2_O_2_ treatment caused a massive shift in TSC22D4 signal to a higher molecular weight, indicating the introduction of new PTM events (Fig. 2D). Previous studies also predicted that TSC22D4 might contain several phosphorylation sites by in silico analysis [9]. To address, whether it is the phosphorylation events that cause the shifts in TSC22D4 western blots, we performed an *in vitro* λ phosphatase assay on immunoprecipitated Flag-TSC22D4. Upon λ phosphatase treatment, the Flag-TSC22D4 signal indeed shifted to a lower molecular weight, confirming that TSC22D4 undergoes phosphorylation (Fig. 4C, compare lanes 4-5 and 10-11).

**Figure 4.**
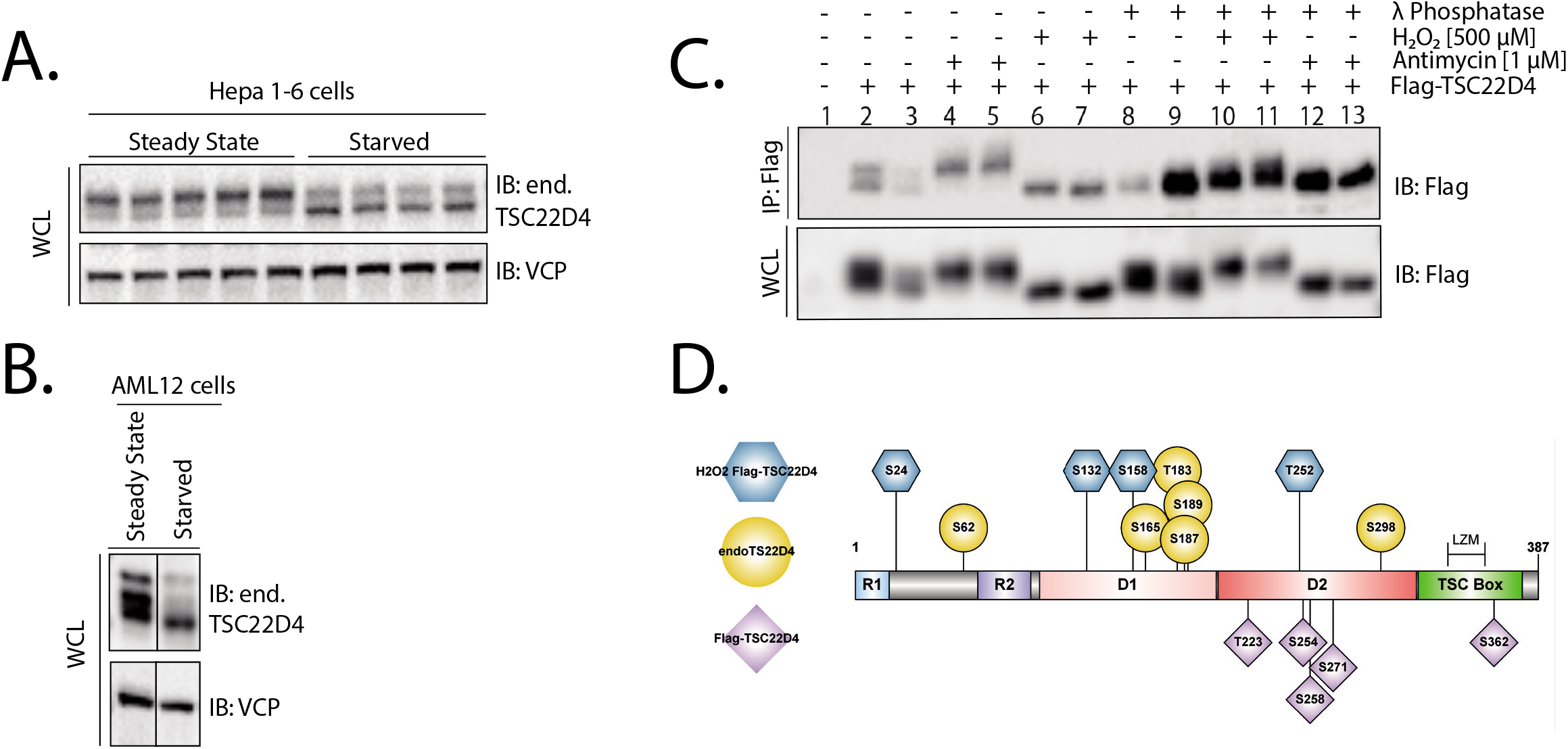
TSC22D2 undergoes phosphorylation. **A and B.** Hepa 1-6 cells (A) and AML12 (B) cells were grown under steady state conditions and were starved or not starved in the absence of glucose and serum for 16 h. Cells were lysed the next day and the whole cell lysates (WCL) were immunoblotted (IB) with indicated antibodies. **C.** Hepa 1-6 cells were transiently transfected with Flag-TSC22D4. 24 h posstransfection cells were treated with H_2_O_2_ or antimycin for 1 h prior to lysis. Flag-TSC22D4 was immunoprecipitated (IP) with anti-Flag affinity gel. IPed Flag-TSC22D4 was subjected to in vitro λ phosphatase treatment. The IPs and WCL were immunoblotted with indicated antibodies. **D.** TSC22D4 phosphorylation sites (p-sites) identified by LC-MS/MS. Yellow: p-sites that are identified on endogenous TSC22D4 in Hepa 1-6 cells (S165) or in mouse livers (S62, T183, S187, S189, S298). P-sites that are identified on transiently transfected Flag-TSC22D4 without or with H_2_O_2_ treatment are indicated in purple and blue, respectively. LZM: Leucine Zipper Motif

Interestingly, although oxidative stress and mitochondrial inhibition caused opposite effects on TSC22D4 phosphorylation, they both promoted the TSC22D4-Akt1 interaction, indicating that the general hyper- or hypo-phosphorylation status of TSC22D4 does not necessarily direct the TSC22D4-Akt1 interaction. Instead, these phosphorylation events might act in a specific manner depending on the direct upstream regulators and control TSC22D4 function accordingly. Hence, we decided to identify the exact amino acid residues on TSC22D4 that undergo phosphorylation. By employing liquid chromatography tandem mass spectrometry (LC-MS/MS) we identified 15 novel phosphorylation sites on TSC22D4 either in Hepa 1-6 cells or in in mouse livers which were highly conserved (Supp. Fig. 2 and Fig. 4D). Interestingly, 6 out of 15 phosphorylation sites lie within the TSC22D4 D2 domain which is required and sufficient to maintain TSC22D4-Akt1 interaction (Fig. 4D). Hence in the future it will be extremely critical to address: **a.** Whether these phosphorylation sites also play role in TSC22D4-Akt1 interaction and **b.** If yes, to identify the responsible kinases. Addressing these questions will certainly help us unravel the functional significance of TSC22D4-Akt1 interaction in regulating Akt1 function.

## Discussion

Type 2 diabetes is a chronic and progressive disorder characterized by elevated blood glucose levels. Although a wide range of type 2 diabetes medications exits, they all have their limitations and side effects. Hence, a better understanding of molecular mechanisms that govern the pathogenesis of insulin resistance and type 2 diabetes, will help development of more effective therapies. Recently, we identified that TSC22D4 impairs insulin-induced Akt phosphorylation and exacerbates diabetic hyperglycemia and insulin resistance [7]. The exact molecular mechanisms of TSC22D4 action in metabolic control however remained poorly defined. Here, we discovered that TSC22D4 is a novel Akt1 interacting protein and TSC22D4-Akt1 interaction responds to metabolic signals such glucose and insulin. Intriguingly, TSC22D4 interacts specifically with Akt1, but not with Akt2 (data not shown), which may explain isoform specific functions that are governed only by Akt1 but not by Akt2 or vice versa. Our data also pointed out to an interesting interplay between glucose and insulin stimulations in controlling TSC22D4-Akt1 interaction. Insulin weakened the TSC22D4-Akt1 interaction only in the presence of glucose and was not sufficient to modulate TSC22D4-Akt1 interaction on its own (Fig. 1C). This single observation leads to at least two intriguing conclusions:

1. Glucose availability renders the TSC22D4-Akt1 interaction insulin-sensitive via yet-to-be identified mechanisms.
2. Akt S473 phosphorylation does not (or is not sufficient to) blunt TSC22D4-Akt1 interaction in the absence of glucose.

Unlike in glucose starved cells, insulin injection to the fasted mice did abolish the TSC22D4-Akt1 interaction, indicating that even low amounts of blood glucose levels during fasting is sufficient to prime the insulin-induced dissociation of TSC22D4-Akt1 interaction (Fig. 1E). Together with our previous data showing that hepatic TSC22D4 regulates hyperglycemia and insulin sensitivity, we are tempted to hypothesize that TSC22D4 controls glucose metabolism via its interaction with Akt1.

TSC22D4 contains a long stretch of intrinsically disordered region. Recent evidence indicates that proteins with intrinsically disordered regions act as critical environmental sensors [19]. Particularly, multiple posttranslational modifications in the intrinsically disordered regions enable protein-protein interactions and promote crosstalk between different signaling pathways Our observations indeed support the notion that TSC22D4 might also acts as an environmental sensor because TSC22D4-Akt1 interaction and TSC22D4 phosphorylation also respond to intra- or extra-cellular cues. Interestingly while both mitochondrial inhibition and oxidative stress promoted the TSC22D4-Akt1 interaction, each had opposite effects on TSC22D4 phosphorylation, indicating that these phosphorylation events act in a rather specific manner to regulate the TSC22D4-Akt1 interaction. In other words, the general hypo- or hyper-phosphorylation status of TSC22D4 does not necessarily control the degree of TSC22D4-Akt1 interaction. Interestingly 6 out of the 15 novel phosphorylation sites that we identified on TSC22D4 lie within the D2 domain, which is both required and sufficient to maintain the TSC22D4-Akt1 interaction. Hence, it will be very intriguing to identify, in the future, whether these phosphorylations play a direct role in maintaining TSC22D4-Akt1 interaction and if so, it will be critical to generate the phospho-specific antibodies and identify the responsible kinases and signaling pathways.

TSC22D4 also needs its TSC box to interact with Akt1 (Fig. 3B). TSC box contains a leucine zipper motif which enables its homodimerization or potentially its heterodimerization with other TSC22 family members. Hence, rather than the TSC box per se, TSC22D4 might actually require to homodimerize or heterodimerize to interact with Akt1. Indeed, in the *in vitro* binding assays we performed with purified recombinant TSC22D4 and Akt1 proteins, TSC22D4 and Akt1 did not interact with each other (data not shown), suggesting that TSC22D4 does not interact with Akt1 directly but rather involves other proteins. Considering the fact that other TSC22 family members and Akt share many common functions, it will be interesting, in the future, also to address the possibility of protein-protein interactions between other TSC22 family members and Akt isoforms.

Based on our findings in this study, we postulate a working model, whereby TSC22D4 acts as a sensor of metabolic and stress signals to interact with Akt1 and regulate its function. Nevertheless, current evidence we present in this study is not adequate to conclude whether the TSC22D4-Akt1 interaction is beneficial or harmful for metabolic health. It might as well have a dual function in a condition-dependent manner. It will be interesting to unravel, in the future, the direct implications of TSC22D4-Akt1 interaction in pathogenesis of several diseases such as metabolic syndrome that involve insulin resistance and type 2 diabetes as well as carcinogenesis, in which dysregulated Akt function plays a key role.

## Materials and Methods

### 1. Animal experiments

Male albumin-cre mice crossed to the TSC22D4 floxed/floxed line (C57Bl/6N background) [20] were housed at room temperature with 12h light-dark cycle on control diet (Research diets, New Brunswick, NJ, USA, D12450B), 60% high fat diet (Research diets, D12492i) with Sucrose Containing Drinking Water (42 g/l) or chow diet (10% energy from fat, Research Diets D12450Ji, USA) for weeks indicated for each study. Mice had *ad libitum* access to food and water and were weighed regularly and inspected daily for general health. All experiments were performed in accordance with the European Union directives and the German animal welfare act (Tierschutzgesetz) and were approved by local authorities (Regierungspräsidium Karlsruhe, license #G117/18 or #G116/18).

For adenovirus injections, 2×10^9^ plaque-forming units (p.f.u.) per recombinant virus were administered via tail vein injection. In each experiment, 4-12 animals received identical treatments and were analyzed under fasted, random fed or fed conditions as indicated. Organs including liver, epididymal and inguinal fat pads, and gastrocnemius muscles were collected after specific time periods, weighed, snap-frozen and used for further analysis.

AAVs were injected via tail vein at a dose of at 1×10^1o^ vector genome/mouse. Before sacrificing post 12-week of AAV injection, mice were starved for 6 h and followed by i.p. injection with 1.5U insulin per kg body weight for 10 min. Organs including liver, epididymal fat pads, kidney, pancreas and gastrocnemius muscles were collected after specific time periods, weighed, snap frozen and used for further analysis. In each animal experiment, mice were randomly assigned to each group.

During animal group allocation animals in each group had similar mean and median body weight as well as fasting blood glucose levels.

### 2. Recombinant viruses

Adenoviruses expressing the Flag-TSC22D4 cDNA under the control of the CMV promoter was cloned using the BLOCK-iT Adenoviral expression system (Invitrogen). Viruses were purified by the cesium chloride method and dialysed against phosphate-buffered-saline buffer containing 10% glycerol before animal injection, as described [21].

For generating AAVs with cDNAs of TSC22D4 alleles (WT or ΔD2) were amplified with the following primer pairs followed by Nhe1 and Xba1 digestion and subcloning into pdsAAV-LP1 plasmid: TSC22D4-AAV-F with NheI: gatgctagcgtgtgctggaattctg, TSC22D4-AAV-R with XbaI: gcatctagactcgagtcagatggaggg. The successful clones sequenced for confirmation with the following primers: pdsAAV-TSC-F: ctgataggcacctattggtc, pdsAAV-TSC-R: ccacaactagaatgcagtg. Once the sequence is confirmed, the plasmids were purified by using Qiagen Megaprep kit (#12381) according to the manufacturer’s instructions and sent to Vigene Biosciences (Maryland, USA) for AAV generation, purification, and titration.

### 3. Primary hepatocyte isolation and treatment

Primary hepatocytes were isolated as previously described [22, 23]. To investigate insulin signaling pathway, primary hepatocytes were seeded to collagen coated 6-well plates (1×10^6^ cells per well) in William’s medium (Pan Biotec, #P04-29510) supplemented with 2 mM L-Glutamine, 100 nM Dexamethasone, 10% Fetal Bovine Serum (FBS, life technologies, #10270-106) and 1% penicillin-streptomycin (PenStrep, Life Technologies, #15140122). After 4 hours, the cells were washed 2 times with phosphate buffered saline (PBS) to get rid of the cells which do not attach to the surface. After 48 hours, the cells were collected in 180 *μl of* 1.5x sample buffer, boiled at 95° for 5 minutes and kept at −20°C until use for western blotting.

### 4. Cell lines and cell culture

Alpha mouse liver 12 (AML12, ATCC, CRL-2254), U2 osteosarcoma (U2OS, ATCC, HTB-96), Hepatoma 1-6 (Hepa1-6) (ATCC, CRL-1830) and human embryonic kidney 293A (HEK293A) cell lines were used. AML12 cells were maintained in DMEM:F12 (ATCC, #302006) medium supplemented with 10% Fetal Bovine Serum (FBS, life technologies, #10270-106), insulin-transferrin-selenium mix (ITS mix, Gibco, #41400045) and 40 ng/ml dexamethasone (Sigma, #D8893). U2OS (wild type and AMPK α1/α2 double knock out) cells were a kind gift from Reuben Shaw (Salk Institute for Biological Studies, San Diego, CA, USA) and were maintained in DMEM supplemented with 10% FBS. Hepa1-6 and HEK293A cells were maintained in Dulbecco’s modified Eagle’s medium (DMEM, Gibco, #11995) supplemented with 10% heat inactivated FBS and 50 U/ml penicillin-streptomycin (PenStrep, Life Technologies, #15140122). HEK293A cells (Thermo Fischer Scientific, #R70507) cells were used to generate adenoviruses. All the cell lines were maintained at humidified 37°C incubator containing 5% CO_2_.

### 5. Plasmid DNA transfection

pcDNA3-HA-Akt1 plasmid was a kind gift from Dr. Diane Fingar (University of Michigan, Ann Arbor, USA). Expression vectors of wild type (WT)-TSC22D4 and deletion mutants of TSC22D4 were created using pcDNA3 plasmid backbone. For deletion mutants, site directed mutagenesis experiments were performed with standard PCR-based methods (NEB, #E0554S) using corresponding primer sets:

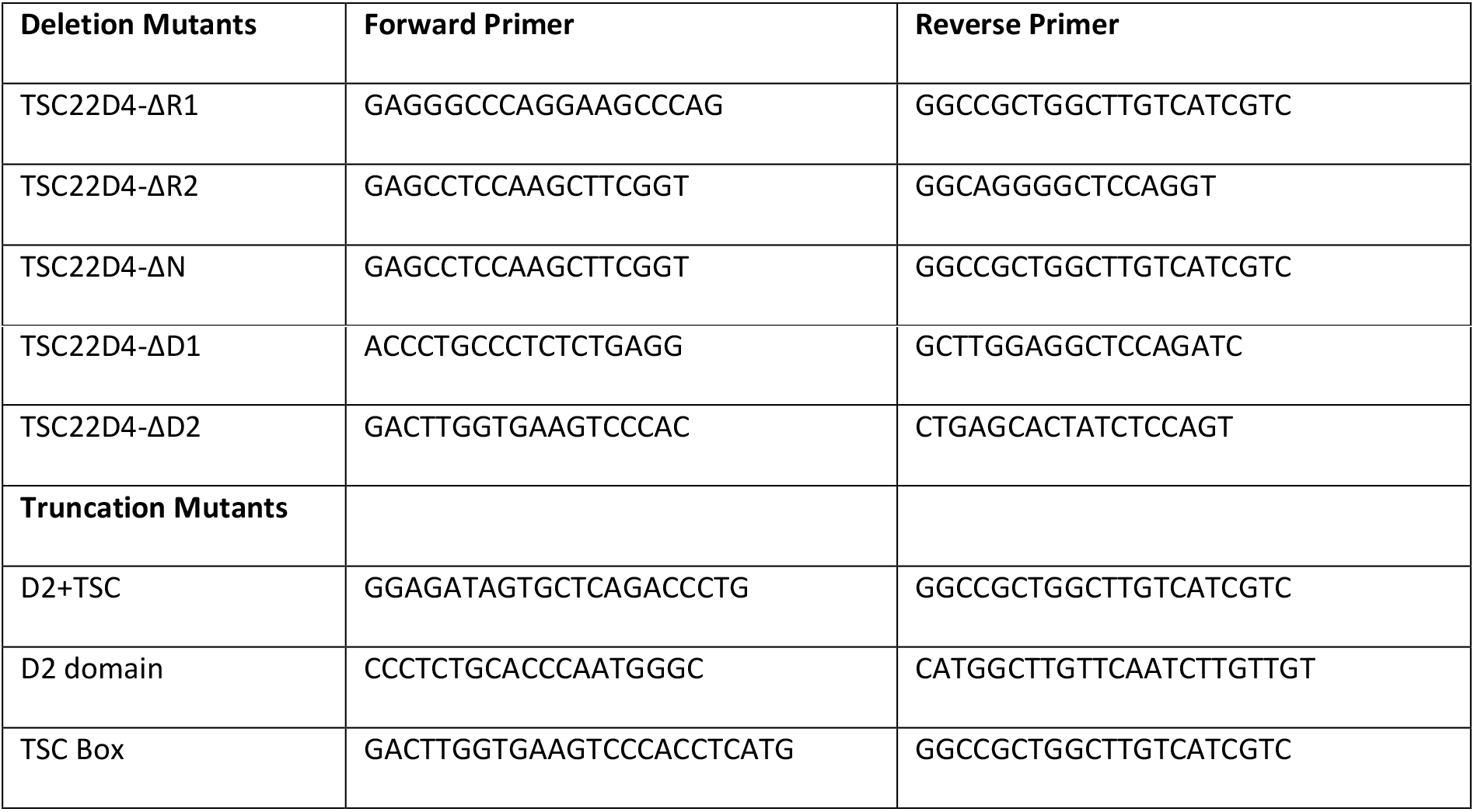

Plasmids expressing TSC22D4 alleles and Akt1 were transiently co-transfected to Hepa1-6 cells using Lipofectamine 2000 (Invitrogen, #11668019) according to the manufacturer’s instructions. For starvation experiments: 30-hour post transfection, the cells were starved in no glucose containing RPMI medium in the absence of FBS (Gibco, #11879020) overnight and the next day cells were pre-stimulated with glucose [20 mM] for 30 minutes and with insulin [100 nM] for an additional 30 minutes followed by cell lysis. For experiments that did not involve starvation, cells were lysed 48-hour post transfection. For U2OS cells, the same transfection protocol was followed except that the cells were starved for 6 hours.

### 6. Identification of TSC22D4 Phosphorylation Sites

#### a. In cultured cells

##### Tryptic digestion and phosphopeptide enrichment

Total cell lysates equivalent to 300 μg of protein per condition were used for a chloroform-methanol precipitation for sample clean up. Followed by a standard in-solution digestion using Trypsin in a 1:100 ratio. The resulting peptides were enriched for phosphorylation using TiO_2_ beads. Prior to MS analysis, the peptides were resuspended in 0.1% TFA/0.25% hexafluoroisopropanol.

##### Mass spectrometry

Tryptic peptides mixtures were separated using a nano Acquity UPLC system (Waters GmbH, Eschborn, Germany). Peptides were trapped on a nano Acquity C18 column (180 μm x 20 mm, particle size 5 μm) (Waters GmbH, Eschborn, Germany). The liquid chromatography separation was performed on a C18 column (BEH 130 C18 100 μm x 100 mm, particle size 1.7 μm (Waters GmbH, Eschborn, Germany)) with a flow rate of 400 nL /min. For all gel slices samples, the chromatography was carried out using a 3h gradient of solvent A (98.9% water, 1% acetonitrile, 0.1 % formic acid) and solvent B (99.9% acetonitrile and 0.1% formic acid) in the following sequence: from 0 to 4% B in 1 min, from 4 to 30% B in 140 min, from 30 to 45% B in 15 min, from 45 to 90% B in 5 min, 10 min at 90% B, from 90 to 0% B in 0.1 min, and 9.9 min at 0% B. The nanoUPLC system was coupled online to an LTQ Orbitrap XL mass spectrometer (Thermo Scientific, Bremen, Germany). The mass spectrometer was operated in the sensitive mode with the following parameters: capillary voltage 2400V; capillary temperature 200°C, normalized collision energy 35 V, activation time 30000 ms. Data were acquired by scan cycles of one FTMS scan with a resolution of 60000 at m/z 400 and a range from 370 to 2000 m/z in parallel with six MS/MS scans in the ion trap of the most abundant precursor ions. The mgf-files generated by Xcalibur software (Thermo Scientific, Bremen, Germany) were used for database searches with the MASCOT search engine (version 2.4.1, Matrix Science, London, UK) against the SwissProt database (Taxonomy Mus musculus; version 2016_05). Peptide mass tolerance for database searches was set to 5 ppm and fragment mass tolerance to 0.4 Da. Carbamidomethylation of C was set as fixed modification. Variable modifications included oxidation of M, deamidation of NQ and phosphorylation STY. One missed cleavage site in case of incomplete trypsin hydrolysis was allowed. Furthermore, proteins were considered as identified if more than one unique peptide had an individual ion score exceeding the MASCOT identity threshold (ion score cut-off of 22-23). Identification under the applied search parameters refers to False Discovery Rate (FDR) < 3,5% and a match probability of p<0.05, where p is the probability that the observed match is a random event.

#### b. In mouse livers

Protein digest and desalting as well as phosphopeptide enrichment were performed as previously described [24]. Briefly, protein lysates were reduced with dithiothreitol (Merck) and cysteine residues were alkylated with iodoacetamide (Merck). In-solution protein digestion was performed using Lys-C (Wako Chemicals, Neuss) and trypsin (Promega). For analysis of the input samples, a 10 μg aliquot of the digest was desalted with Pierce C18 spin columns (Thermo Fisher Scientific) according to the manufacturer’s instructions, with 1 μg subjected to analysis by LC-MS/MS. The remaining part of the digest was acidified with TFA prior to desalting by solid-phase extraction using Sep-Pak C18 cartridges (Sep-Pak tC18, 1 cc Vac (100 mg), Waters). Phosphopeptides were enriched by titanium dioxide (TiO_2_) chromatography as described [24].

LC-MS/MS analysis was performed on a Q-Exactive HF mass spectrometer (Thermo Scientific) online coupled to an Ultimate 3000 nano-RSLC (Dionex). Peptides were automatically loaded on a C18 trap column (300 μm inner diameter (ID) × 5 mm, Acclaim PepMap100 C18, 5 μm, 100 Å, LC Packings) at 30μl/min flow rate prior to C18 reversed phase chromatography on the analytical column (nanoEase MZ HSS T3 Column, 100Å, 1.8 μm, 75 μm x 250 mm, Waters) at 250nl/min flow rate in a 95 minutes non-linear acetonitrile gradient from 3 to 40% in 0.1% formic acid. Profile precursor spectra from 300 to 1500 m/z were recorded at 60000 resolution with an automatic gain control (AGC) target of 3e6 and a maximum injection time of 50 ms. TOP10 fragment spectra of charges 2 to 7 were recorded at 15000 resolution with an AGC target of 1e5, a maximum injection time of 150 ms for phospho-measurements and 50 ms for measurements of unenriched samples, an isolation window of 1.6 m/z, a normalized collision energy of 27 and a dynamic exclusion of 30 seconds.

Mass spectrometric raw data was processed separately for the phospho-enriched and non-enriched fraction of samples with Progenesis QI for proteomics (Version 3.0, Nonlinear Dynamics, Waters) to perform label-free quantification with pairwise normalization as described in [25, 26] and Proteome Discoverer (Version 2.1.1.21, Thermo Fisher Scientific) using the Sequest HT search engine and the ptmRS node for the confident localization of phosphosites [24].

### 7. Cell or tissue lysis and co-immunoprecipitation (Co-IP)

For co-IP experiments, cells and frozen liver tissues were lysed in ice-cold lysis CHAPS-buffer containing 10 mM KPO4 (pH 7.2), 1 mM EDTA, 5 mM EGTA, 10 mM MgCl2, 50 mM β-Glycerophosphate and 0.3% CHAPS supplemented with Complete Protease Inhibitor Cocktail (Roche, #11836145001) and PhosSTOP EASYpack phosphatase inhibitor (Roche, #4906837001). Lysed cells or liver tissue were centrifuged at 12.000 rpm for 5 minutes at 4°C and supernatants were collected. Protein concentrations were measured with Bradford (BioRad, #5000006) or BCA assay (Pierce, #23227). For immunoprecipitation of transfected cell lysates, anti-Flag affinity gel (Sigma, #A2220) or HA conjugated agarose beads (Sigma, #A2095) were used. Prior to use, the beads were washed with lysis buffer via centrifugation at 4.000 rpm for 2 minutes. For immunoprecipitation, cell lysates were incubated with the beads for 2 hours at 4°C on a rotator. The beads were washed 3 times with ice-cold co-IP lysis buffer with centrifugation steps in between. 2x laemmli buffer was added to the samples which then were boiled at 95°C for 5 minutes.

Liver lysates of WT mice were prepared to analyze endogenous interaction of TSC22D4-Akt1. Rabbit IgG control [1 mg/ml] (CST, #2729S) or TSC22D4 antibody (Pineda, home-made antibody, 1mg/ml) were incubated with liver lysates at 4°C overnight on a rotating rack. Next day, protein A/G beads (Santa Cruz, #sc-2003) were conjugated to protein-antibody complex with 2 hours of incubation at 4°C on a rotating rack. The lysates were washed 3 times with lysis buffer and added 70 *μl of* 2x laemmli sample buffer.

### 8. Western blot analysis

Protein extracts and immunoprecipitates were run on 10% (BioRad, #4561036) or any kD (BioRad, #4569036) precast protein gels in sodium dodecyl sulfate (SDS) buffer followed by electrophoretic transfer of proteins to nitrocellulose membrane (BioRad, #2895). After blocking in 5% nonfat dried milk diluted in tris-buffered saline and 1% tween 20 (TBS-T) for 1 hour at RT, the membranes were incubated with primary antibodies specific for TSC22D4 (home-made, by PINEDA Antibody Service, Flag-HRP (#A8592, Sigma), HA (#2999, CST), Myc (#2040, CST), p-Akt (S473) (#4060, CST), p-Akt (T308) (#13038, CST), Akt (#9272 ,CST), p-GSK3 beta (S9) (#05-643, Upstate), GSK3 beta (#05-412, Upstate), p-Foxo1 (S256) (#9461, CST), Foxo1 (#2880, CST), p-p70S6K1 (#9234, CST), p70S6K1 (#9209, CST), pS6 (#2271, 2215), S6 (#2217, CST), pAMPK α T172 (#2531, CST), AMPK α (#2532, CST), VCP (ab11433, Abcam). Primary antibodies were incubated in 3% BSA overnight at 4°C on a rocker. After 3 times wash with TBS-T each for 10 minutes, membranes were incubated with HRP-conjugated secondary antibodies targeting rabbit or mouse IgG for 1 hour at RT. To avoid background noise in endogenous co-IP experiments, HRP-conjugated veriblot (Abcam, #131366) was used as secondary antibody. Immunoblots were developed using ECL western blotting substrate (Sigma, #GERPN2209 or Amersham, #GERPN2236). ChemiDoc Imaging System (BioRad) was used to detect and analyze chemiluminescence signals.

### 9. *In vitro* λ phosphatase assay

Immunoprecipitated Flag-TSC22D4 was subjected to λ phosphatase assay as previously described [27]. Following Flag-TSC22D4 immunoprecipitation, beads were washed three times in IP lysis buffer followed by two additional washes in ST buffer (50 mM Tris-HCl, pH 7.2; 150 mM NaCl). Beads were then resuspended in 1x phosphatase buffer (50 mM Tris-HCl, pH 7.5, 100 mM NaCl, 2 mM dithiothreitol, 0.1 mM EGTA, 0.01% Brij 35) that contained 2 mM MnCl2. Samples were incubated at 30°C for 30 min in the absence or presence of λ -phosphatase (Invitrogen), and reactions were terminated by adding EDTA, pH 8.0, to a 50 mM final concentration and sample buffer to a 1x final concentration.

### 10. Software and Data analysis

Adobe Illustrator and IBS (Illustrator for biological sequences) programs were used to create figures and cartoons of deletion mutants.

### 11. Image Editing

For some figures, irrelevant lanes were removed from western blot images. Such modifications were indicated with a thin black line on western blot figures.

## Acknowledgments

We would like to thank Diane Fingar (University of Michigan, Ann Arbor, USA), Reuben Shaw ( Salk Institute for Biological Studies, San Diego, CA, USA), Thomas Fleming (Heidelberg University Hospital, Germany), Dominic Helm (DKFZ, Heidelberg) and Bilgehan Ibibik (Bilkent University, Ankara, Turkey) for their support during the preparation of this manuscript.

---

**Supplementary Figure 1.**
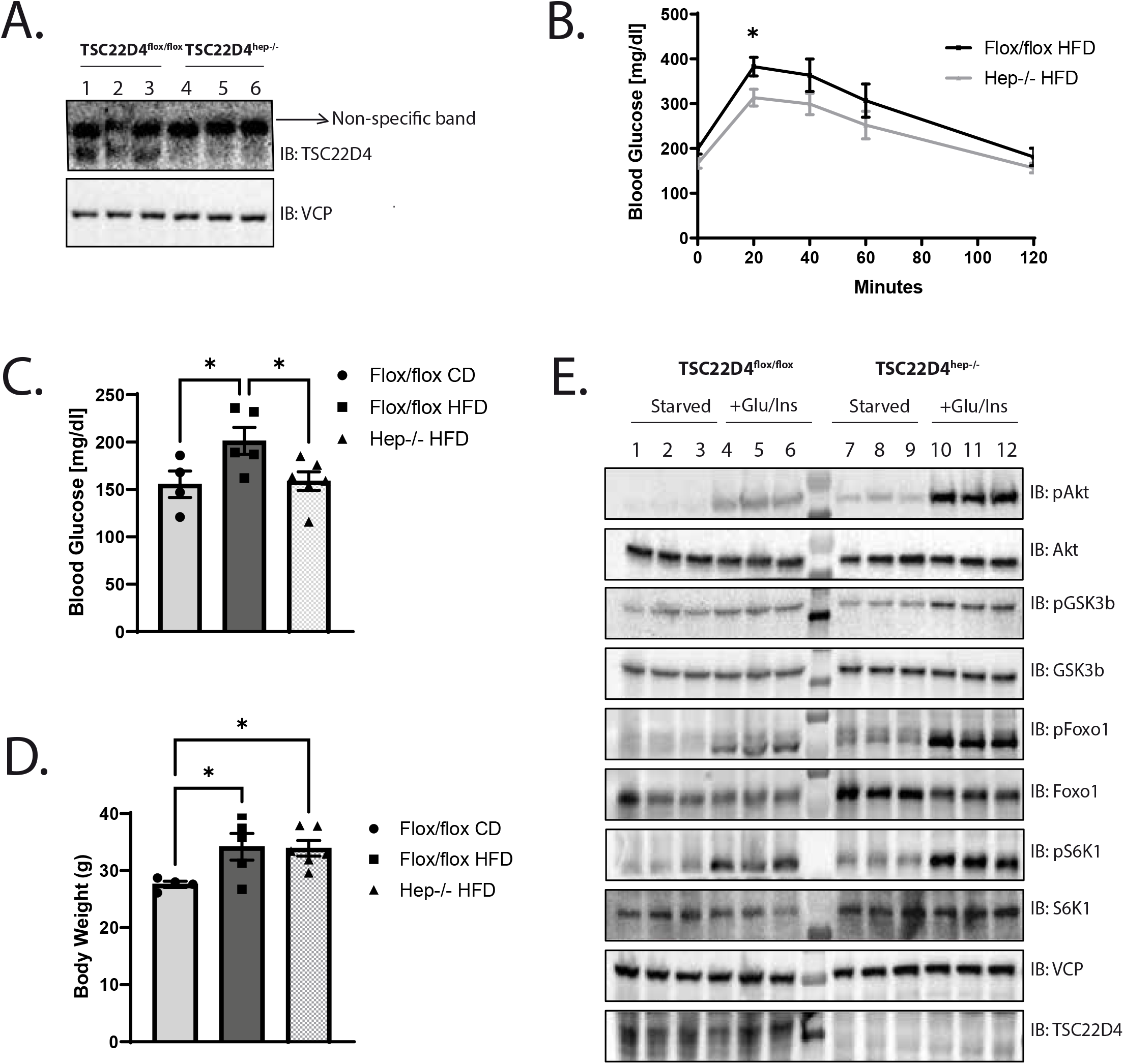
Hepatocyte specific TSC22D4 knockout (TSC22D4^hep-/-^) mice show imroved glucose handling. **A.** Western blot analysis of the mouse liver lysates from TSC22D4^fox/fox^ control mice or TSC22D4^hep-/-^ mice immunoblotted with indicated antibodies. **B.** Intraperitoneal glucose tolerance test (i.p. GTT) performed with mice as in A subjected to high fat diet (HFD) for 6 weeks. Mice were fasted for 6 h of followed by glucose solution injection at a dose of 1g/kg mouse weight. n=5-6 per group. Statistical analysis: 2-way ANOVA with repeated measures and Fisher’s LSD test. **C.** Fasting blood glucose levels of TSC22D4^fox/fox^ and TSC22D4^hep-/-^ mice on 9 weeks of control diet (CD) or HFD challenge. Statistical analysis 2-way ANOVA and Fisher’s LSD test. **D.** Body weight of mice as in C. Statistical analysis 2-way ANOVA and Dunnet’s multiple comparisons test. **E.** Western blot analysis of primary hepatocytes of TSC22D4^fox/fox^ control mice and TSC22D4^hep-/-^ mice in starved or glucose and insulin stimulated conditions immunoblotted (IB) with indicated antibodies.

**Supplementary Figure 2.**
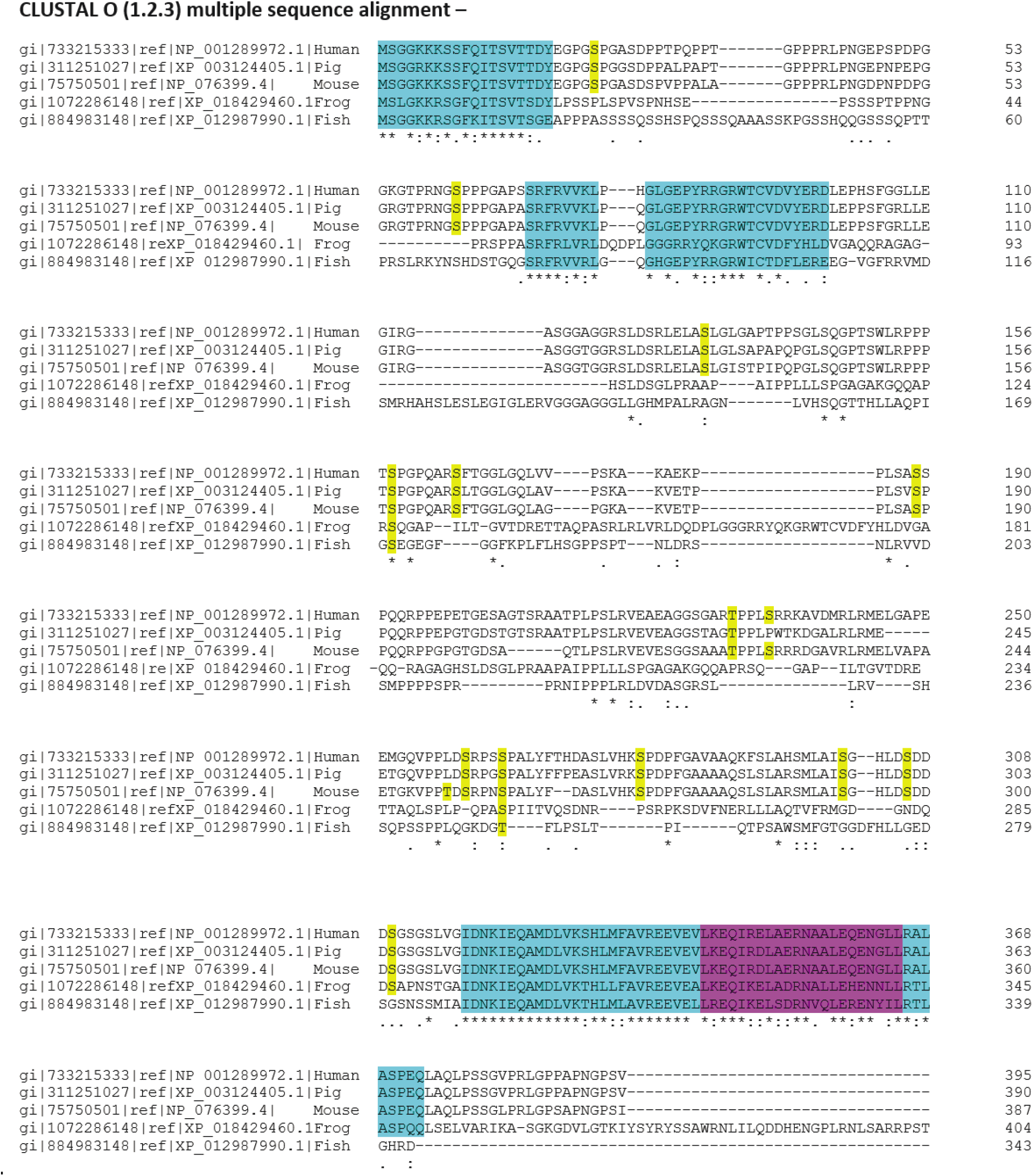
CLUSTAL Omega multiple sequence alignment analysis of TSC22D4. Highlighted in blue: Evolutionarily conserved regions on TSC22D4 primary amino acid sequence. Highlighted in pink: Leucine Zipper Motif Highlighted in yellow: Novel phosphorylation sites on TSC22D4 identified via LC/MS-MS

**Supplementary Figure 3.**
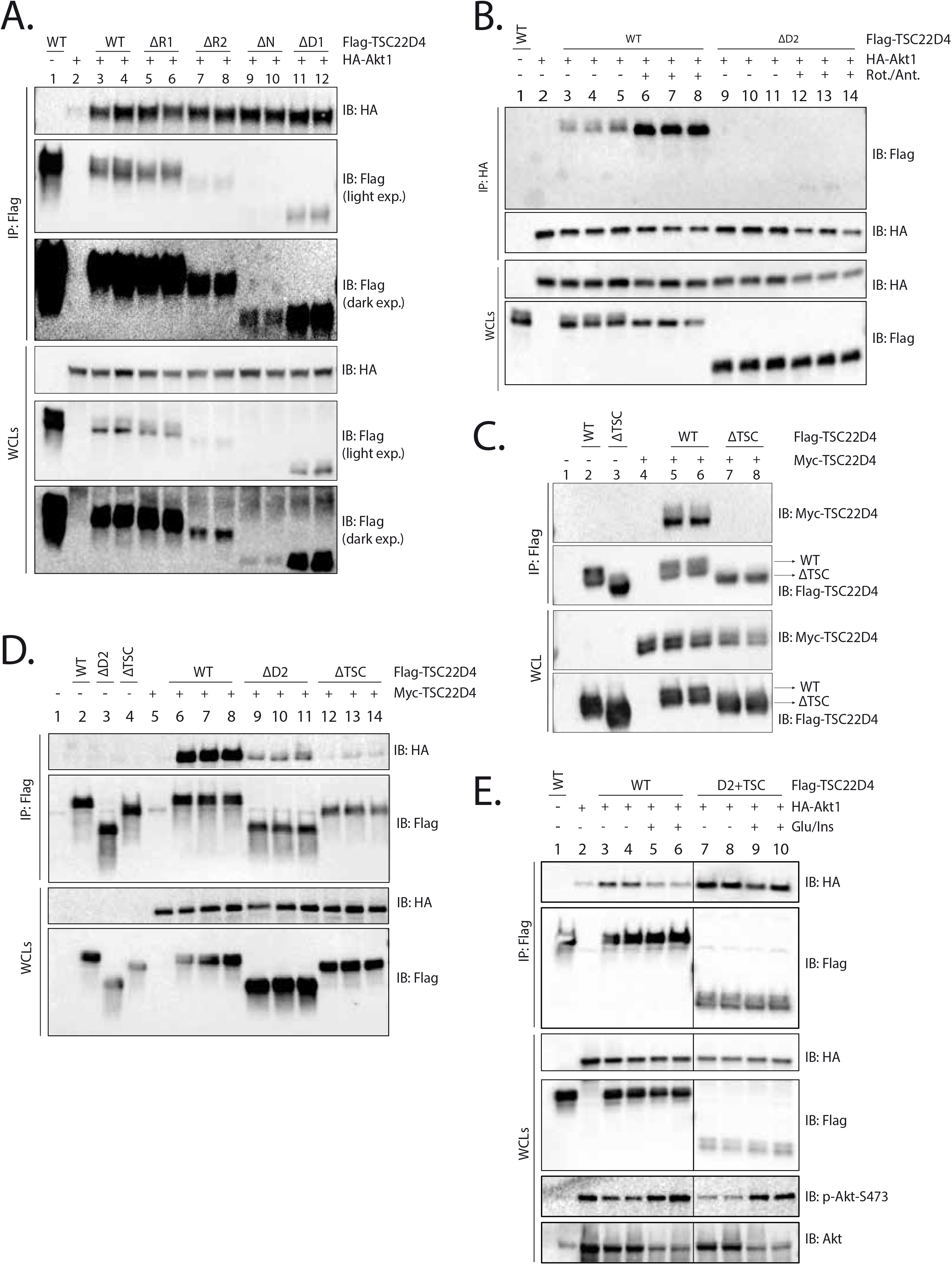
Mapping the Akt1 interaction domains on TSC22D4. **A**. Hepa1-6 cells were transiently co-transfected with Flag-TSC22D4-WT (2,5 μg), -ΔR1 (2,5 μg), -ΔR2 (2,5 μg), -ΔN (2,5 μg) or -ΔD1 (2,5 μg) deletion mutants and HA-Akt1 (2,5 μg). Flag-TSC22D4 was immunopecipitated (IP) with anti-Flag affinity gel and the IPs and WCLs were immun-blotted (IB) with indicated antibodies. **B.** Hepa1-6 cells were transiently co-transfected with Flag-TSC22D4-WT (2,5 μg) or -Δ D2 (2,5 μg) and HA-Akt1 (2,5 μg). 30 h post transfection, cells were serum and glucose starved overnight and the next day, they were treated without or with Rotenone/Antimycin [1 μM] for 1 h prior to cell lysis. IPs were performed with anti-HA antibody. IPs and WCLs were immunblotted (IB) with indicated antibodies. **C.** Hepa1-6 cells were transiently co-transfected with Flag-TSC22D4-WT (2,5 μg) or -ΔTSC (2,5 μg) and Myc-TSC22D4-WT (2,5 μg). Flag-TSC22D4 alleles were immunoprecipitated with anti-Flag affinity gel (IP) and WCLs were immunoblotted (IB) with indicated antibodies. **D.** Hepa1-6 cells were transiently co-transfected with Flag-TSC22D4-WT (2,5 μg), -ΔD2 (2,5 μg) or -ΔTSC (2,5 μg) and Myc-TSC22D4-WT (2,5μg). Flag IPs and WCLs were immunoblotted (IB) with indicated antibodies. **E.** Hepa1-6 cells were transiently co-transfected with Flag-tagged WT-TSC22D4 (2,5 μg) or D2+TSC (2,5 μg) and HA-Akt1 (2,5 μg). 30 h post transfection, cells were serum and glucose starved overnight and the next day, they were stimulated without or with glucose [20 mM] for 30 minutes and insulin [100 nM] for another 3o minutes. Flag IPs and WCLs were immunblotted (IB) with indicated antibodies.

